# Where Tabular Foundation Models Falter on Genetic Data: Datasets That Expose and Provide a Path to Address the Gap

**DOI:** 10.64898/2026.09.25.754520

**Authors:** Anirban Das, Yan Cui

## Abstract

Tabular foundation models are used as off-the-shelf predictors for heterogeneous tabular tasks, but it remains unclear how they will perform on real genotype-to-phenotype tabular datasets, which carry unique challenges. One such challenge is ancestry-dependent non-stationarity in allele effect sizes (the effects of genetic variation on disease risk): the predictive contribution of a given genetic variant may change significantly across the ancestry spectrum. This is especially consequential for patients from ancestry groups that are underrepresented in existing datasets, because models that are unaware of this non-stationarity are more likely to perform poorly for them. The battery of synthetic tasks used to pretrain tabular foundation models does not capture this structure, and we show that it leads to systematic performance degradation. Using controlled hierarchical Gaussian-process stress tests, we demonstrate that both off-the-shelf TabICL and TabPFN are robust when ancestry dispersion in the data is low, but as dispersion grows, the latent ancestry-dependent non-stationarity of effect sizes becomes a first-order problem, and the predictive performance of both models deteriorates. We confirm the same pattern on All of Us (AoU) (a large biobank containing whole-genome sequencing and electronic health record (EHR) data spanning the ancestry spectrum) across an extensive panel of cancer phenotypes evaluated with the two leading tabular foundation models: TabICL and TabPFN. Holding the in-context training-set size, phenotype, and feature set fixed, ancestry-specific in-context tables, which are less dispersed in ancestry space, consistently outperform size-matched meta-ancestry tables, isolating ancestry dispersion as the driver of degradation. To address the failure without changing the base architecture, we construct two synthetic task families that explicitly encode ancestry-dependent effect drift and instruction-tune an off-the-shelf TabICL model on tasks sampled across both families. On held-out AoU evaluations covering cancer and respiratory disease phenotypes, the tuned model yields more stable performance across ancestry-distance bins and is especially strong in the bins farthest from the center of the in-context exemplars in ancestry space, that is, on subjects whose ancestry is most underrepresented among the provided exemplars. The paper contributes an evaluation protocol, a failure analysis, and two synthetic task families targeted at ancestry-dependent non-stationarity in precision-medicine tabular modeling.

## 1 Introduction

Tabular foundation models such as TabPFN (Hollmann et al., 2025) and TabICL (Qu et al., 2025) are attractive for biomedical prediction (Das and Cui, 2026) because they can be deployed with limited task-specific optimization. This promise is especially relevant for genotype-to-phenotype prediction, where labeled datasets are often modest for underrepresented ancestry groups even when the parent biobank is large (Sirugo et al., 2019; Gurdasani et al., 2019; Mills and Rahal, 2020; Bien et al., 2019). In this setting, each subject is represented by allele dosages at selected single-nucleotide polymorphism (SNP) loci together with ancestry-sensitive covariates derived from genome-wide variation, and the learning problem is to predict a binary phenotype such as a cancer or respiratory outcome.

A well-documented difficulty is that genotype-to-phenotype relationships are often not stationary across ancestry space. If *x*_*j*_ denotes the dosage of SNP *j* and *a* denotes a subject’s location in a continuous ancestry representation, then a natural model is

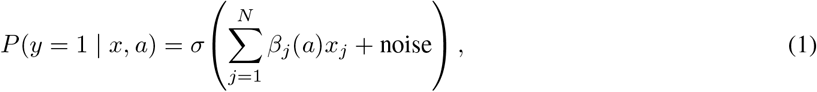

where stationarity corresponds to replacing *β*_*j*_(*a*) by a constant coefficient. Recent work on polygenic prediction shows that accuracy varies continuously with genetic distance from the training population (Ding et al., 2023; Kachuri et al., 2024), and large multi-ancestry studies report population-dependent effect sizes and genetic architectures (Tcheandjieu et al., 2022; Wang et al., 2023). Yet the synthetic tasks typically used to pretrain tabular foundation models are built around broad causal diversity rather than ancestry-dependent effect-size drift.

This paper explores and defines a strategy to address this gap through synthetic data generation and task construction. Synthetic data play a major role in aligning models to targeted issues (Das et al., 2025, 2026). The generators create tables with ancestry-dependent effect drift; the task setup then determines which rows are provided as in-context exemplars and which rows are held out for prediction. The tasks are arranged for the generalization problem: robust prediction for individuals from data-disadvantaged backgrounds who lie far from the effective training center in ancestry space. Our starting point is the empirical question: when real genotype-to-phenotype data violate stationarity, how do off-the-shelf tabular foundation models fail, and can synthetic task design repair the failure without changing the base architecture? The main contribution is not a new general-purpose architecture, but an evaluation protocol together with synthetic dataset families and task constructions that expose a specific failure mode. We show that instruction tuning on our dataset families improves performance on a wide range of real genotype-to-phenotype examples.

A primary barrier to equitable genomic prediction is that biobank resources are highly imbalanced across ancestry groups, with far more training subjects from European-ancestry populations than from data-disadvantaged groups (Sirugo et al., 2019; Gurdasani et al., 2019; Mills and Rahal, 2020; Bien et al., 2019). That imbalance makes non-stationarity especially consequential: when a target subject from an underrepresented group is far removed from the European-ancestry training region, the relevant SNP effect sizes can differ substantially from those estimated near the training center (Ding et al., 2023; Kachuri et al., 2024; Tcheandjieu et al., 2022; Wang et al., 2023). Models that are blind to ancestry-dependent non-stationarity therefore mis-estimate variant effects precisely where equitable generalization matters most.

Contributions: First, we use controlled hierarchical Gaussian-process (HGP) tasks to show that off-the-shelf TabICL and TabPFN perform stably under near-stationary effect sizes but can collapse once training data become more ancestrally dispersed and effect sizes change across ancestry space. Second, we transfer that logic to AoU cancer phenotypes using matched ancestry-specific and meta-ancestry training sets, showing that increased dispersion predicts reduced off-the-shelf performance in real data. Third, we introduce two synthetic task families for alignment: an HGP family that offers direct control of effect-size smoothness and a PRS-CSx-inspired family (Ruan et al., 2022). Fourth, we show that instruction tuning an off-the-shelf TabICL model on mixed HGP and PRS-CSx tasks improves robustness across ancestry-distance bins for both cancer and respiratory disease outcomes in AoU. The AoU program is uniquely suited to serve as the real-data testbed: it provides whole-genome sequencing data from participants spanning the ancestry continuum, with participants recruited from diverse populations (All of Us Research Program, 2024).

## 2 Background

One related approach, from Thomas et al. (2024), is to retrieve target-relevant examples and fine-tune the tabular model on that local subset before prediction. This addresses distributional heterogeneity through local adaptation, but changes one of the main attractions of tabular foundation models for biomedical deployment: their zero-shot, no-retraining use at evaluation time. Applied to genotype-to-phenotype tables, such a method would introduce a new optimization step for each local region or sample.

In the ancestry setting, local retrieval followed by local fine-tuning is also conceptually close to what Gao and Cui (2020) call an independent learning scheme, in which data from different ethnic groups are used separately to train separate models. Prior work in supervised biomedical prediction shows that completely separating groups can be inferior to transfer-learning strategies, because not all alleles exhibit the same degree of effect-size non-stationarity across ancestry groups (Gao and Cui, 2020). Some variant effects are highly ancestry-dependent, while others are comparatively stable and can be learned more accurately by borrowing information across the ancestry space. Both families of synthetic datasets constructed in this paper deliberately encode this property: allele effect sizes are not assumed to be equally non-stationary across alleles.

A second related line studies tabular distribution shift over time. Drift-Resilient TabPFN models temporal non-stationarity by constructing tasks in which future samples must be predicted from past samples, with task generation based on structural causal models that drift over time (Helli et al., 2024). This setting is qualitatively different from the ancestry-dependent effect-size drift considered here. Temporal drift imposes an ordering and a past-to-future prediction problem, and the drift occurs over a one-dimensional space.

Finally, we use the term *instruction tuning* for the act of taking an off-the-shelf TabICL model and further training it on our synthetic task families. We borrow the term from language-model instruction tuning (Wei et al., 2021), but our usage is not identical. In FLAN-style instruction tuning, training examples include task descriptions and in-context exemplars, and evaluation can use a task description alone. In our tabular setting, both tuning and evaluation are mediated only through in-context exemplars; the “instructions” are implicit in the structure of the synthetic tabular tasks rather than written prompts. We acknowledge this abuse of notation.

## 3 Why Non-Stationarity Matters for Genotype-to-Phenotype Tables

The failure mode studied here is not simply class imbalance or lack of examples. It arises from the interaction between two properties. The first is *effect-size non-stationarity*: the same allele can have different predictive value for subjects at different locations in ancestry space (Ding et al., 2023; Tcheandjieu et al., 2022; Wang et al., 2023). Crucially, if all subjects of interest have very similar ancestral backgrounds (the cohort has very little dispersion in ancestry space), non-stationarity of allele effect sizes across the broader ancestry space has little effect when studying this local cohort. The problem arises only when a second property also prevails: *training-set dispersion*. Once a training set (for tabular foundation models these are the exemplars) covers a broad ancestry region, a single set of context examples or learned summary statistics is forced to average over locally different genotype-to-phenotype mappings. That combined effect is especially important for data-disadvantaged communities, because the subjects farthest from the effective training center are precisely those for whom a stationarity assumption is most harmful (Peterson et al., 2019; Kachuri et al., 2024).

This perspective suggests a concrete prediction. If two training sets have the same size but one is more dispersed in ancestry space, then under non-stationarity the more dispersed set should yield worse prediction, especially for models unable to address or recognize non-stationarity. Sections 4 and 5 test this prediction first in synthetic data (where we control both factors) and then in AoU (where we control data dispersion) for tabular foundation models.

## 4 Controlled Stress Tests Reveal the Failure Mode

To stress-test the failure mode in a controlled setting, we create families of tasks. We begin with a simple Gaussian-process family.

### A single task: step-by-step data generation

Fix a configuration (*l, S*), where *S* is a sampling distribution over ancestry space. Each individual *i* is assigned ancestry coordinates *a*_*i*_ ~ *S ⊂* ℝ^*A*^, where *A* is the dimension of the ancestry representation, together with dosage features *G*_*ij*_ *∈* {0, 1, 2} for *j* = 1, …, *N*. The phenotype label follows a, Bernoulli model with logistic link, 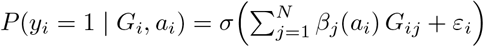 where each effect-size function *β*_*j*_(*·*) is sampled as a Gaussian process (Seeger, 2004) over ancestry space with RBF-style correlation

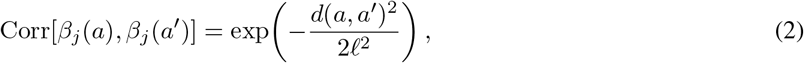

where *d*(*a, a’*) is Euclidean distance in ancestry space. In this illustrative GP construction all effect functions are non-stationary; large *ℓ* makes the sampled processes nearly constant across ancestry, while small *ℓ* induces sharp local heterogeneity. The resulting *M* samples are organized into a table with ancestry coordinates, dosage features, and phenotype label. The table is split into *train* rows (labeled in-context exemplars) and *test* rows (labels masked); the tabular foundation model is asked to complete the masked labels in the standard in-context evaluation setup.

By varying (*ℓ, S*), we obtain a *family of tasks*. Varying *ℓ* controls effect-size non-stationarity, and varying *S* controls the dispersion of samples over ancestry space.

### The HGP family used in this paper

The actual task family used for the stress tests is the *Hierarchical GP* (HGP) extension of this simple GP construction. HGP adds biologically motivated details on top of the GP modeling scheme; full details are provided in Appendix C.

Figure 1 shows the resulting failure pattern for off-the-shelf TabICL and TabPFN on these tasks. Each point corresponds to a different task with different effect-size non-stationarity and data dispersion. Under weak non-stationarity, both models remain largely stable as dataset dispersion increases. Under moderate non-stationarity, the decline is gradual. Under strong non-stationarity, however, both models can move from strong performance on homogeneous datasets to near-random behavior once the support set spans a wider ancestry region. The sign and magnitude of the effect depend on the smoothness of the underlying effect functions; this is the signature one would expect if pretrained tabular models lack alignment to ancestry-dependent effect drift.

**Figure 1:**
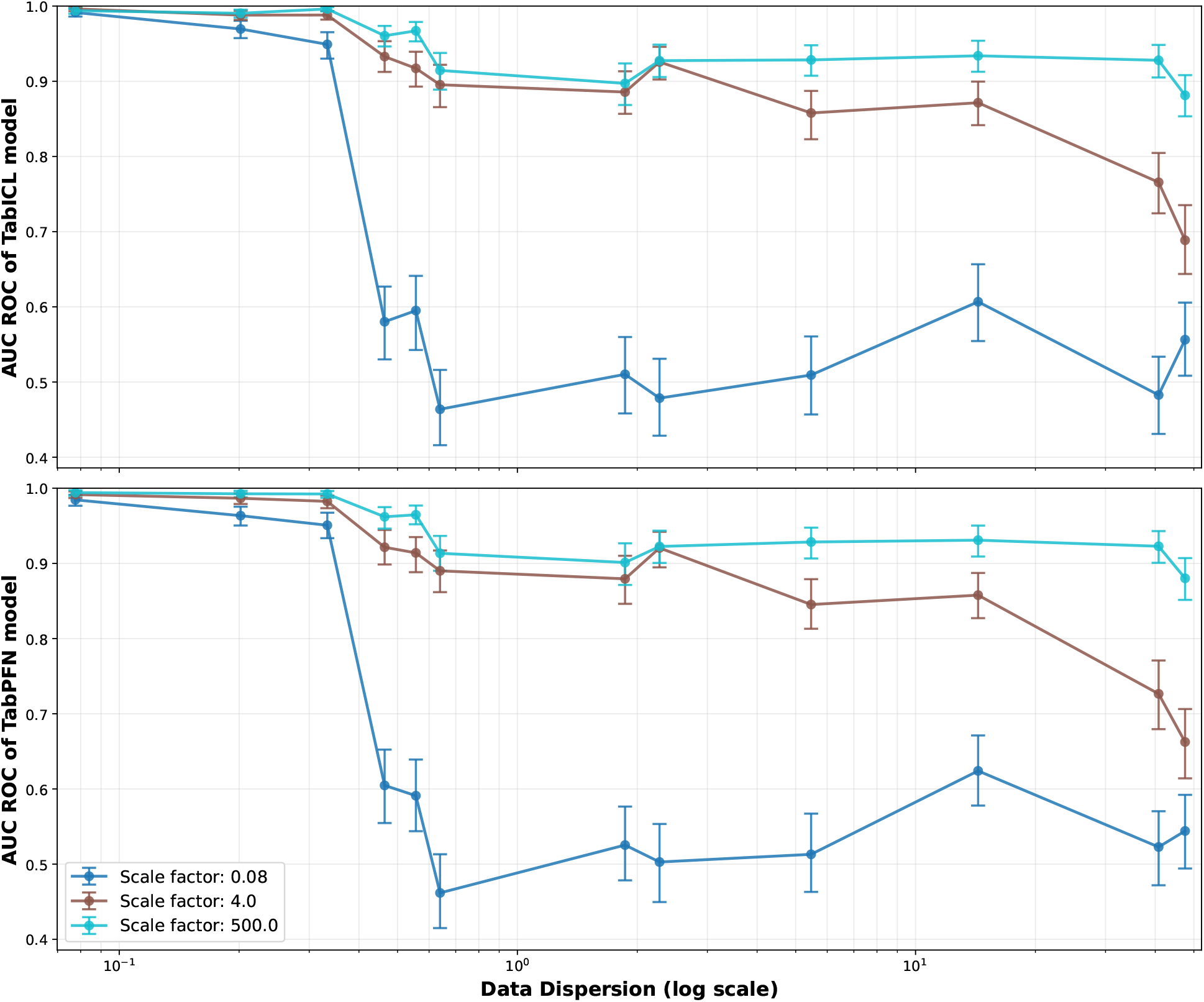
Off-the-shelf TabICL and TabPFN on controlled HGP stress tests. Both models remain stable under weak non-stationarity but degrade sharply as training dispersion (log scale) increases under strong local drift. Curves correspond to three effect-function length-scale factors (0.08, 4.0, and 500.0); error bars show confidence intervals.

## 5 The Same Pattern Appears in Real AoU Cancer Prediction Tasks

We next ask whether the synthetic prediction above appears in real genotype-to-phenotype tables. Modern biobanks store genome-wide genotypes at millions of assayed or imputed SNP loci, but downstream phenotype prediction typically works on a phenotype-specific subset selected through GWAS and linkage-disequilibrium pruning. In our AoU pipeline, each table contains allele dosages for the top 50 phenotype-associated variants, a 16-dimensional continuous ancestry coordinate supplied by the AoU genomic release (All of Us Research Program Genomics Investigators, 2024), and a binary phenotype label. We use the AoU controlled-tier resource (All of Us Research Program Genomics Investigators, 2024) together with the All by All GWAS release built on version 7 participants. Across the six ancestry groups AFR (African), AMR (Admixed American), EAS (East Asian), EUR (European), MID (Middle Eastern), and SAS (South Asian), we retain the 7 cancer phenotypes with at least 10k cases in version 7 and focus on the five data-disadvantaged target groups AFR, AMR, EAS, MID, and SAS. GWAS-based feature selection is performed using all available version 7 participants, and the in-context labeled exemplars are also drawn from version 7. Held-out evaluation is performed only on version 8 samples, which are not used for GWAS, feature selection, or exemplar construction.

The real-data comparison is designed to isolate dispersion while holding sample size fixed. For each phenotype-target ancestry combination with at least 500 available version 7 training samples from the target ancestry, we construct a matched pair of in-context tables. The ancestry-specific table uses complete version 7 training examples from that target ancestry only as the labeled exemplars, and its incomplete rows come from the corresponding version 8 test table for that same target ancestry. The meta-ancestry table uses complete version 7 training examples pooled across all ancestries, then subsampled so that the number of labeled exemplars exactly matches the ancestry-specific table; its incomplete rows come from the version 8 meta-ancestry test cohort for the same phenotype. Each comparison therefore fixes the phenotype and the number of in-context exemplars, while contrasting an ancestry-specific cohort with a more dispersed meta-ancestry cohort. The standard in-context evaluation regime used throughout the paper for AoU data is: no gradient updates are performed on the target AoU dataset before evaluation, the models are evaluated off the shelf, and the training table serves as the set of in-context exemplars.

In Figure 2 we observe that across 16 matched comparisons, the more dispersed meta-ancestry training set yields lower AUC in 13 cases, and 11 of those pairs have non-overlapping 60% bootstrap confidence intervals. Repeating the same matched-dispersion experiment with TabPFN version 6.3.2 yields the same qualitative pattern: in 12 of 16 phenotype-target ancestry combinations, the model performs better with the ancestry-specific context table, which has milder ancestry dispersion, than with the size-matched meta-ancestry context table. The full TabPFN results are provided in Appendix Figure B.1. Because the training size is matched inside each pair, the result cannot be explained by the usual “more data beats less data” argument. Instead, it is consistent with the synthetic diagnosis from Section 4: once effect sizes drift over ancestry space, increasing dispersion without aligning the model to that drift can reduce performance in these matched real-data comparisons.

**Figure 2:**
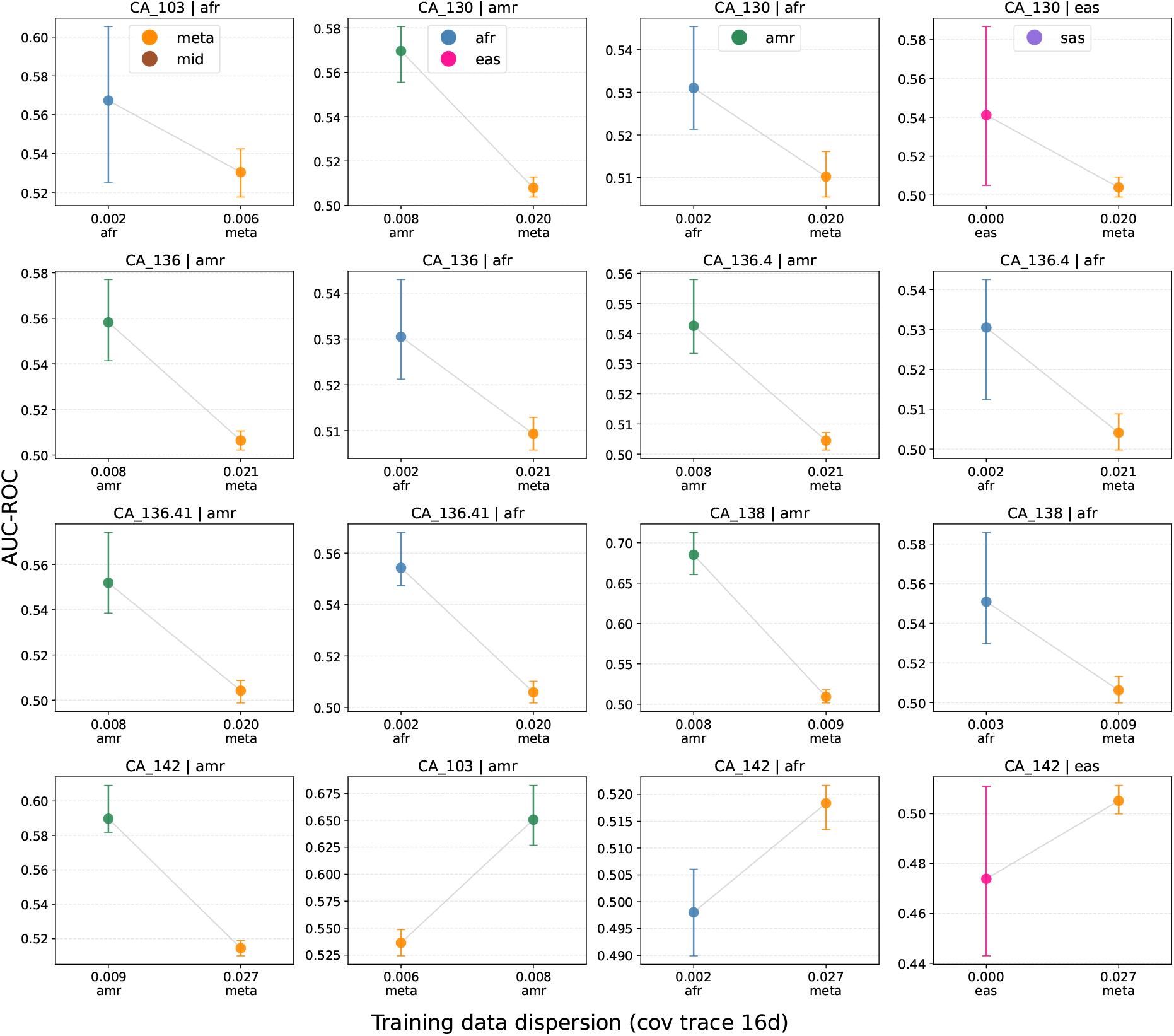
Real-data evidence of the same failure mode in AoU cancer phenotypes. Each panel fixes a phenotype and target ancestry for the ancestry-specific arm, then compares off-the-shelf TabICL using an ancestry-specific exemplar set against a size-matched meta-ancestry exemplar set. In both arms, the horizontal axis reports training-set dispersion in ancestry space and the vertical axis reports AUC on the corresponding evaluation cohort. Only phenotype codes are shown in this figure; full phenotype names are listed in Appendix Table B.1.

**Remarks**. The real-data comparisons are subject to hidden confounders that are absent from the synthetic experiments, and hence results are expected to be noisier. Known confounders are: disease heritability can differ across ancestry groups, linkage disequilibrium structure can differ between the target-ancestry and meta-ancestry cohorts, and the SNP set is discovered from a GWAS performed on the meta-ancestry cohort, with the same loci then reused for the target ancestry subjects (Kachuri et al., 2024). This last factor can bias the setup in favor of the meta-ancestry cohort, making it easier for the meta-ancestry arm to achieve higher AUC-ROC for any model. That bias makes our finding conservative; the more dispersed meta-ancestry tables still underperform in most matched comparisons. A second point to keep in mind is that we use only genetic features. In real clinical prediction, non-genetic features such as age, sex, and laboratory measurements carry substantial predictive signal. We omit those variables here because our goal is to isolate non-stationarity of genetic effect sizes across ancestry space. This omission likely lowers absolute accuracy, but it does not undermine the comparative conclusions. Finally, because all features are noisy genetic features and the target-ancestry evaluation cohorts often contain limited numbers of version 8 test samples, the performance metrics are themselves noisy. For this reason, Figure 2 and Appendix Figure B.1 use 60% bootstrap confidence intervals, corresponding to 20% bootstrap mass in each tail; when these intervals do not overlap, the bootstrap summary indicates at least a 60% chance that the arm with the higher point estimate is truly higher under the resampling distribution.

## 6 Two Synthetic Task Families for Alignment

The failure analysis above motivates a targeted alignment strategy: instruction tune on synthetic tasks with ancestry-dependent effect drift, where a *task family* is simply a parameterized generator and each hyperparameter draw defines one task. We use two complementary task families.

### HGP family

The same HGP generator used in Section 4; see Appendix C for the mathematical description of a single task and the family parameterization. The HGP family combines three ingredients: Gaussian-process effect-function variation across the ancestry continuum, cluster-specific allele-frequency perturbations that produce marginal feature-distribution shifts, and imbalanced cluster sampling that produces uneven sample density across ancestry.

### PRS-CSx family (Ruan et al., 2022)

This family models the same three mechanisms as HGP—ancestry-varying feature-target relationships, marginal feature-distribution shifts, and uneven local sample density—but does so through a parameterization that is closer to biological population-genetic structure. It starts from a continuous-shrinkage prior over SNP effects (a global-local Gamma-Gamma hierarchy that produces sparse heavy-tailed effects), introduces cluster-specific allele frequencies through a Balding-Nichols construction, and couples populations through per-SNP cross-population genetic correlation. Cluster-level effects are interpolated using per-SNP bandwidths. Relative to HGP, this family adds sparse effects concentrated on a small number of causal loci, explicit coupling between effect magnitudes and ancestry-specific allele frequencies, and cluster-specific heritability structure. Full mathematical details of both families are given in Appendices C and D.

### Instruction-tuning protocol

We initialize from the off-the-shelf TabICL checkpoint (version 0.1.4) and retrain on tasks sampled uniformly from the mixed family. For each task, synthetic tabular data are first generated and then converted into an evaluation task with labeled in-context exemplars (70%–90% of rows) and masked test rows. The model attends over the exemplars and predicts the masked labels; loss is cross-entropy, accumulated across many tables per gradient update. Tasks have a deliberate design choice: the majority cluster is oversampled in the exemplar set, while the test set is more evenly spread across clusters. This makes the task demand the behavior we need in practice: using a biased exemplar set to predict individuals whose ancestries are underrepresented among those exemplars. We generate 400 training tasks and 60 held-out evaluation tasks.

## 7 Instruction Tuning Improves Distance-Robustness on Real Data

The comparison in this section is between instruction-tuned TabICL and off-the-shelf TabICL (version 0.1.4). None of the models in this section is retrained or fine-tuned on any AoU tabular data: the tuned model is updated only on synthetic HGP and PRS-CSx tasks, and both models are evaluated on AoU data only through the in-context tables. Full optimization details are deferred to Appendix E.

The evaluation design follows the framework of Ding et al. (2023), which evaluates predictive performance as a function of how far each subject lies from the training distribution. Here we measure that distance from the center of the in-context exemplar set. The reasoning is as follows: a model that implicitly assumes stationarity will, during training, learn variant-phenotype relationships for individuals near the center of the training distribution. As test subjects lie progressively further from this center in ancestry space, those learned relationships become increasingly inaccurate because non-stationarity causes the true relationships to diverge from those learned in the training region. Models that cannot account for non-stationarity therefore exhibit greater prediction error for subjects more distant from the training center. If instruction tuning on HGP and PRS-CSx tasks reduces that sensitivity, it should manifest as flatter performance curves across ancestry-distance bins compared with the off-the-shelf model.

To operationalize this, we use genotype data from AoU together with SNP locations identified by the All by All GWAS, restrict in-context exemplars to European-ancestry version 7 subjects, and evaluate on version 8 meta-ancestry test subjects. Among 46 cancer phenotypes available, we retain the 7 with more than 10k EUR cases in version 7; among 53 respiratory-related phenotypes, we retain all 10 with more than 10k EUR cases in version 7. Version 7 is the source of the in-context exemplar set and the GWAS feature-discovery step. We compute each test subject’s Mahalanobis distance (Mahalanobis, 2018) from the center of the version 7 in-context exemplar set in ancestry-PC space, sort subjects by this distance, and divide them into 10 equal-sized bins. The metric is the mean log-probability assigned to the correct class, reported with 80% bootstrap confidence intervals.

Figures 3 and 4 show the outcome for cancer and respiratory disease phenotypes, respectively. In both cases the off-the-shelf model typically loses performance as test subjects move away from the training center. The instruction-tuned model flattens these distance-performance curves. The farthest distance bin favors the tuned model in all 17 phenotypes.

**Figure 3:**
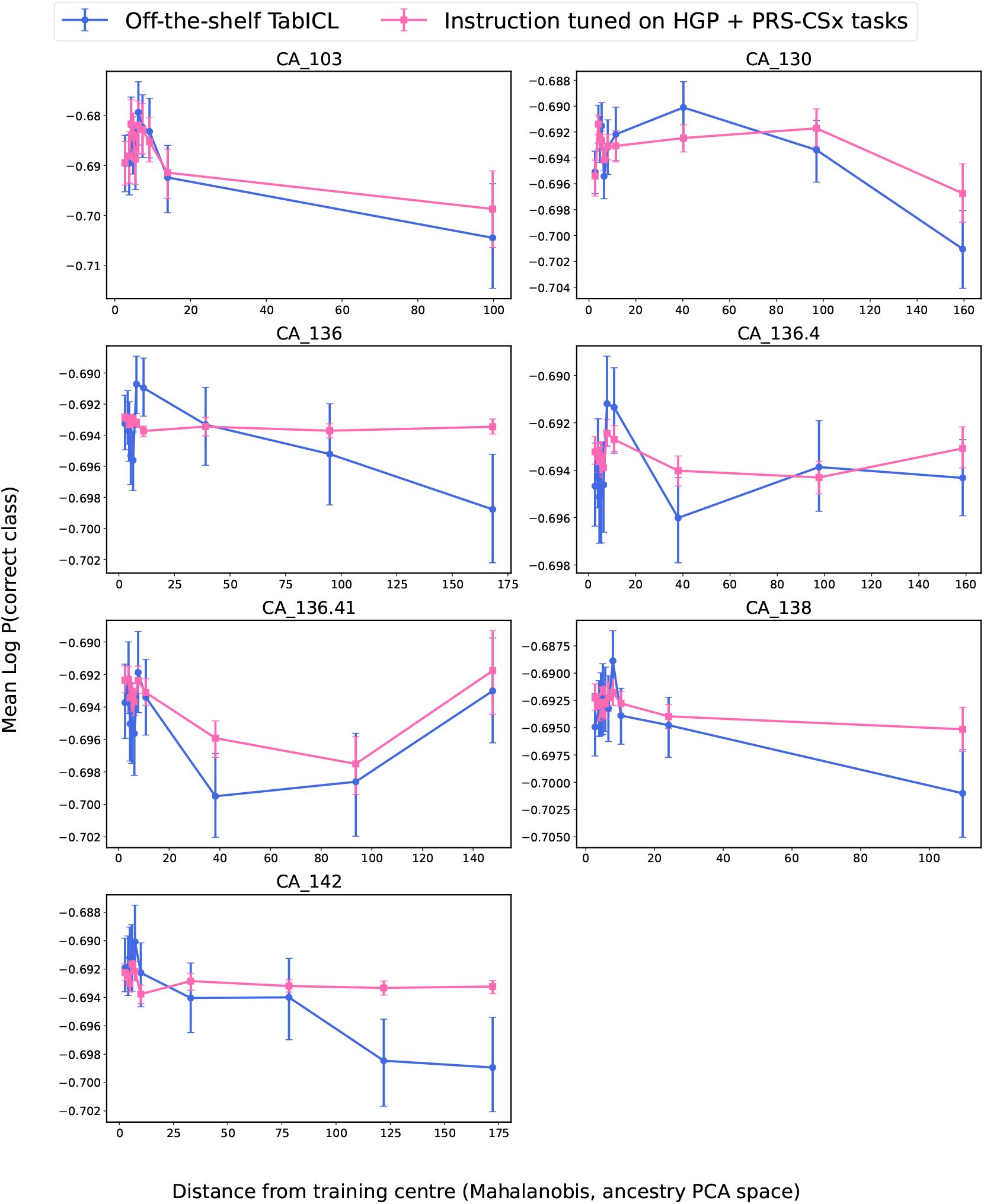
Instruction tuning on mixed HGP and PRS-CSx tasks improves robustness for **cancer phenotypes** in AoU. Mean log-probability of the correct class is plotted versus Mahalanobis distance from the version 7 training center in ancestry-PC space. Error bars are 80% bootstrap confidence intervals. Only phenotype codes are shown in this figure; full phenotype names are in Appendix Table B.1.

**Figure 4:**
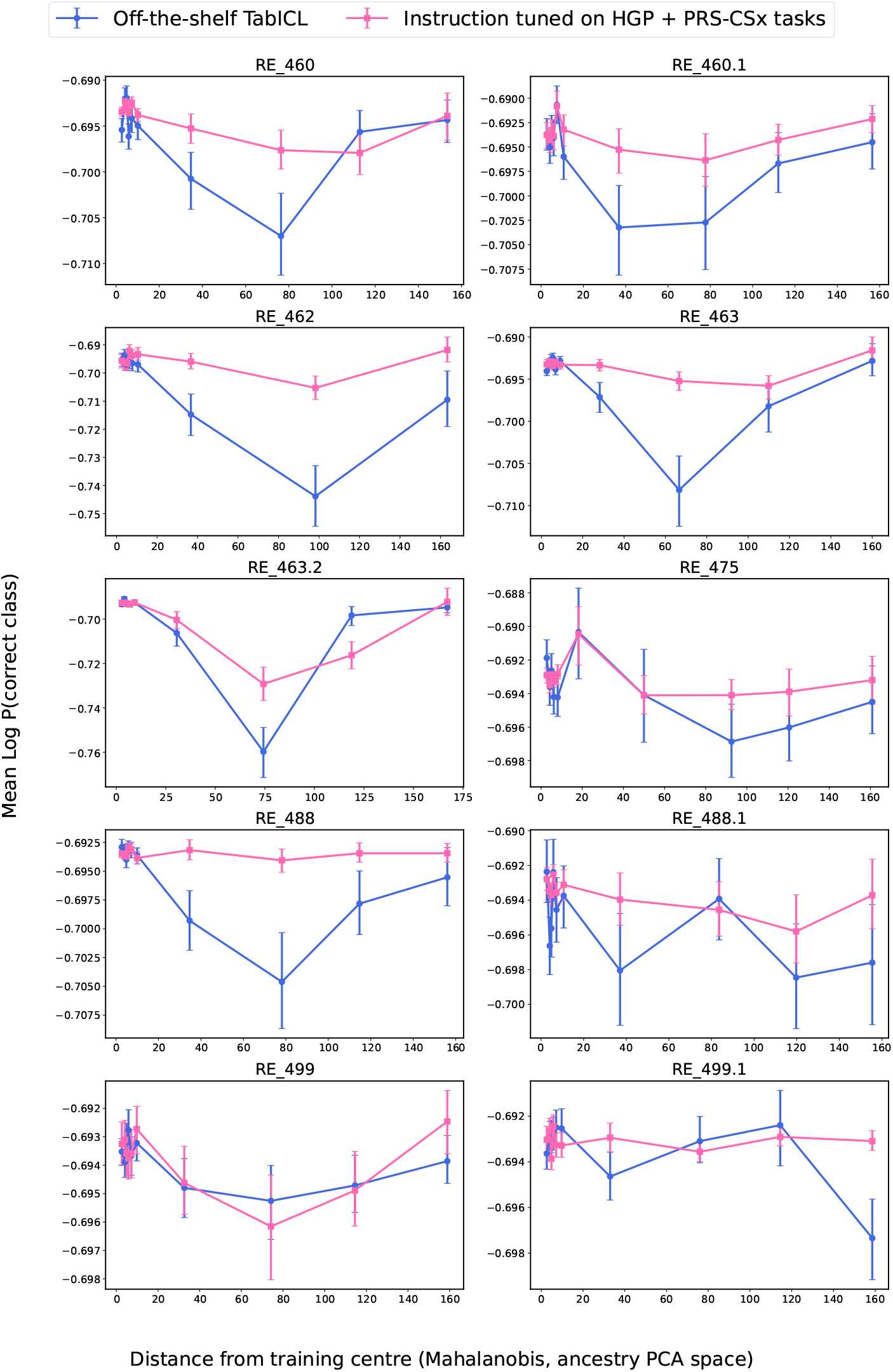
Instruction tuning on mixed HGP and PRS-CSx tasks improves robustness for **respiratory disease pheno-types** in AoU. Same evaluation protocol as Figure 3. Only phenotype codes are shown in this figure; full phenotype names are listed in Appendix Table B.1.

## 8 Discussion and Limitations

This paper is evaluation-centered. The main message is that ancestry-dependent non-stationarity is a reproducible failure mode for off-the-shelf tabular foundation models on genotype-to-phenotype tasks, and that synthetic task design can mitigate the failure in a predictable direction. The evidence is sequential: controlled HGP tasks isolate the mechanism, matched AoU cancer comparisons show the same mechanism in real data, and mixed HGP+PRS-CSx instruction tuning reduces the expected failure on both cancer and respiratory disease phenotypes.

A first practical question is why the large-scale real-data evaluation for instruction tuning centers on TabICL. Here we work in a secondary-analysis regime: GWAS-based SNP discovery is performed on a cohort that is separate from the evaluation cohort. We follow standard doctrine for polygenic risk score analysis in such settings (Choi et al., 2020). Under that design, we impose a conservative threshold of 10k available training samples per phenotype, which in turn requires a model that can handle large in-context tables efficiently. We therefore focus on TabICL, whose transformer architecture scales linearly in the number of rows in the training table and can accommodate AoU tables, some of which have roughly 40k rows. In primary-data settings, by contrast, one can often hold out a subset of samples before GWAS and work with substantially smaller downstream evaluation tables, which may make it practical to compare a broader set of tabular foundation models.

The ancestry-dependent mechanisms modeled in our synthetic task families are ancestry-varying feature-target relationships, marginal feature-distribution shifts, and uneven local sample density. These are not the only obstacles that hinder precision medicine for individuals in data-disadvantaged communities. Another prominent issue is that linkage structure changes across the ancestry continuum, so SNPs discovered in a GWAS cohort can become progressively less informative for individuals farther from that cohort’s center in ancestry space. Our synthetic task families do not directly model this additional ancestry-dependent shift in linkage structure. Measuring how well tabular foundation models can handle linkage-structure change through synthetic data construction remains a necessary step before claiming broader utility for precision-medicine studies.

## AI use statement

In this work, we used generative AI tools to write code that implements methods and to assist with the associated data processing; all AI-generated code was verified manually by the authors. Generative AI tools were not used for any of the other tasks requiring mandatory disclosure under standard AI-use policies for authors: we did not use AI to generate synthetic datasets or to develop the paper’s theoretical models or conceptual frameworks; to formulate mathematical claims or provide critical ingredients for proofs; to assist in writing proofs or to propose or refine hypotheses; to design the research methodology or experiments or to provide feedback on them; or to support qualitative or thematic data analysis or to interpret results.

Among the recommended-disclosure categories, we used generative AI tools (i) to aid in polishing and editing the writing for clarity and readability, (ii) to assist with retrieval and discovery of related work, including identifying relevant literature, (iii) to draft portions of the text of the paper, and (iv) to create and edit software code. We reviewed and edited all AI-assisted writing, and we re-verified every reference cited in this paper directly against primary sources (publisher pages, PubMed, arXiv, and institutional repositories) rather than relying on AI-generated citation content. We take responsibility for the final content of this work, including all text, claims, code, and artifacts produced with the aid of generative AI.

## Acknowledgments

We gratefully acknowledge All of Us participants for their contributions, without whom this research would not have been possible. We also thank the National Institutes of Health’s All of Us Research Program for making available the participant and sample data examined in this study. This study used data from the AoU Research Program’s Controlled Tier Dataset v7 and v8, available to authorized users. Ancestry coordinates and group labels used in this study are provided by the AoU Research Program for research purposes and reflect genetic principal component-based groupings; they should not be interpreted as proxies for race, ethnicity, or social identity. We emphasize the importance of responsible interpretation of ancestry-related findings in genomic research and caution against conflating genetic ancestry with social constructs of race. This work was supported by the US National Cancer Institute grant R01CA262296.

## Reproducibility statement

Code is available at https://anonymous.4open.science/r/Where-Tabular-Foundation-Models-Falter-on-Genetic-Data-Nurips-26-Submission1839/README.md. Full details needed to reproduce the results in this paper are provided in the appendix. The construction of the AoU tabular datasets, including phenotype and feature selection and the ancestry-specific/meta-ancestry matched-dispersion protocol, is described in Appendix B. The complete mathematical specification of the two synthetic task families used for the controlled stress tests and for instruction tuning is given in Appendices C and D. The instruction-tuning protocol, including the loss function, optimizer settings, hyperparameter search grid, and compute resources, is described in Appendix E. Our plan for releasing the synthetic task generators, task manifests, and the tuned model checkpoint is described in Appendix F.

## A Appendix Overview

This appendix records the real-data construction protocol, the mathematical form of the two synthetic task families used for alignment, and the instruction-tuning protocol behind the main-paper results. It is intentionally self-contained so that the synthetic tasks can serve as reusable evaluation resources even without access to the controlled-tier AoU data used in the downstream experiments.

### B Real-Data Construction in AoU

#### B.1 Linking with the All By All GWAS Study

The tabular datasets used in this project are assembled by linking two primary resources from the AoU Research Program (All of Us Research Program Genomics Investigators, 2024): the individual-level controlled-tier short-read whole-genome sequencing (WGS) data and the All By All GWAS release produced by the Broad Institute. The All By All GWAS associated allelic variants across the genome with 2,333 unique phenotypes using stratified analyses across six ancestry groups—AFR, AMR, EAS, EUR, MID, and SAS—as well as a meta-analysis on the unstratified participant pool. All subjects and data for the GWAS came from version 7 of the AoU dataset. This large-scale GWAS analysis was conducted using SAIGE in the Google Cloud environment; by linking to its results we leverage this investment without rerunning the discovery.

For this project, we restrict attention to phenotypes from the phecodeX category for which at least 10,000 cases are observed in the version 7 training population. We focus on cancer and respiratory phenotype families, which yield 7 cancer phenotypes and 10 respiratory phenotypes after applying this threshold. For each phenotype, we retrieve the top GWAS-significant variants from the per-ancestry and meta-ancestry summary statistics, apply linkage-disequilibrium pruning to obtain an approximately independent set, and retain the 50 most significant variants as input features.

#### B.2 Tabular Feature Construction

The end product of the linking pipeline is a set of tabular training and test datasets. Each table contains three components: (1) allele dosage features for the 50 selected variants, (2) the 16-dimensional ancestry principal-component (PC) coordinates provided directly by the AoU genomic release, and (3) a binary phenotype label. The 16-dimensional ancestry PC vector is a continuous summary of genome-wide allele-frequency patterns; it naturally embeds individuals along the genetic ancestry continuum and is the same representation used in the AoU program’s own analyses.

Age and sex matching is performed before the final tabular dataset is frozen wherever needed to avoid confounding by demographic factors unrelated to the genetic signal under study.

#### B.3 Training and Test Splits and In-Context Evaluation Design

We exploit the AoU program’s two-version release structure to enforce a strict temporal separation between feature discovery and final assessment. Feature selection (top GWAS hits, LD pruning) is performed using version 7 participants only. Training tables are drawn from version 7 subjects; test tables are drawn from subjects who enrolled in the program in version 8 and were therefore not available to the upstream GWAS.

For the matched-dispersion analysis of real-data degradation (Section 5 of the main paper), each phenotype–target ancestry comparison uses two training tables with exactly the same sample count: an ancestry-specific table containing only version 7 subjects from the target ancestry group, and a meta-ancestry table containing version 7 subjects from all ancestry groups subsampled to match the ancestry-specific table size. Evaluation is identical for both conditions: training subjects serve as in-context labeled exemplars, while the disease labels of test subjects from the corresponding target-ancestry version 8 table are masked and the tabular foundation model is asked to predict them. No gradient updates are performed at test time; the training table plays the role of the context window. The training-dispersion statistic shown in the main paper is the trace of the ancestry-coordinate covariance matrix of the training subjects.

#### B.4 Additional Matched-Dispersion Results with TabPFN

Figure B.1 repeats the main-paper matched-dispersion analysis using TabPFN version 6.3.2. The pattern is consistent with the TabICL result in the main paper: in 12 of 16 phenotype–target ancestry combinations, the model performs better when the in-context exemplars come from the ancestry-specific version 7 table, which has milder ancestry dispersion, than when they come from the size-matched meta-ancestry version 7 table. As in the main analysis, all evaluation samples are version 8 subjects and no model parameters are updated on AoU tabular data.

For the instruction-tuning evaluation (Section 7 of the main paper), in-context exemplars come from European-ancestry version 7 subjects, while test subjects come from the version 8 meta-ancestry cohort. Test subjects are ordered by Mahalanobis distance from the version 7 training center in ancestry-PC space and split into 10 equal-sized bins. If *a* is a test subject’s ancestry coordinate, *µ* is the mean ancestry vector of the training set, and Σ is the regularized training covariance, then the distance statistic is

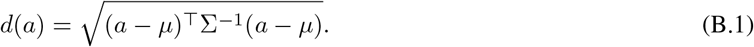

The plotted quantity is the mean log-probability of the correct class inside each bin, with 80% bootstrap confidence intervals.

### C Hierarchical Gaussian-Process Task Family

The HGP family provides direct, interpretable control over ancestry-dependent non-stationarity. It is the same family used for both the main-paper stress tests (Section 4) and the instruction-tuning curriculum. All implementation details are in `make_data_ver3.py` and `test_train_dist_shift.py.`

#### C.1 Population and Cluster Structure

Let *K*_*c*_ denote the number of ancestry clusters. Cluster centers *c*_*k*_ *∈* ℝ*A* are drawn i.i.d. from a standard normal and then scaled so that the mean pairwise inter-cluster distance equals a reference distance *L*_scale1_.

##### Imbalanced cluster assignment

Subjects are assigned to clusters with deliberately unequal proportions to mimic the ancestry imbalance of real biobanks. The *K*_*c*_ clusters are partitioned into ancestry groups by cutpoints; the major group (one cluster) receives weight 1.0, while subsequent groups receive weights 1*/d*_1_ and 1*/d*_2_ (*d*_1_ = 9, *d*_2_ = 12 by default). Approximately (1 −*π*_noise_) ×100% of subjects are allocated according to these weights, and the remaining *π*_noise_ fraction are assigned by sampling cluster indices uniformly at random. After assignment, cluster labels are shuffled.

##### Per-cluster width

A base scale *L*_scale2_ = *ν · L*_scale1_ (default *ν* = 0.004) sets the overall within-cluster spread. Each cluster *k* is assigned a width multiplier *m*_*k*_ drawn from a discrete set (default {0.5, 0.75, 1.0, 1.5}), with the constraint that the majority cluster always satisfies *m*_0_ = 2 × max_*k*≠0_ *m*_*k*_, making it both more populated and spatially wider than any minority cluster.

Individual *i* in cluster *k*_*i*_ has ancestry coordinates drawn as

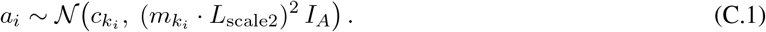

#### C.2 Genotype Generation with Cluster-Specific Allele Frequencies

For SNP *j*, a base minor-allele frequency is drawn as *p*_*j*_ *~* Uniform(*p*_min_, *p*_max_). A cluster-specific perturbation

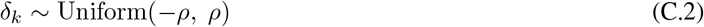

is applied to yield cluster-specific MAFs:

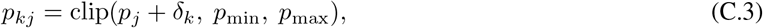

where *ρ* is the cluster MAF offset (default *ρ* = 0.08). The perturbation *δ*_*k*_ is shared across all SNPs for cluster *k*, producing a genome-wide MAF shift between clusters. Dosages are then sampled as

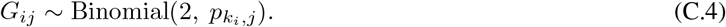

#### C.3 Effect Functions via Gaussian Processes

All *N* SNPs receive ancestry-varying effect functions. Each SNP *j* is independently assigned a lengthscale

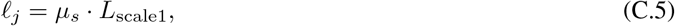

where the scale multiplier *µ*_*s*_ is drawn uniformly from a discrete set (default {100.0, 0.5, 0.008}). These three regimes correspond respectively to near-stationary, moderately non-stationary, and strongly non-stationary effect functions.

The *M* realized effect values for SNP *j* are drawn jointly as

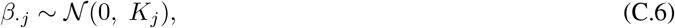

where *K*_*j*_ *∈* ℝ ^*M* ×*M*^ has entries

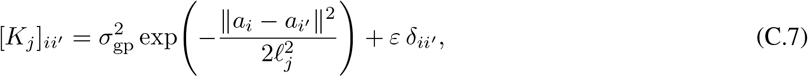

with diagonal jitter *ε >* 0 for numerical stability. A linear drift *m*(*a*) = *a*^*T*^*b* can be added to the GP mean (default *b* = 0).

#### C.4 Phenotype Model

The latent logit for subject *i* is

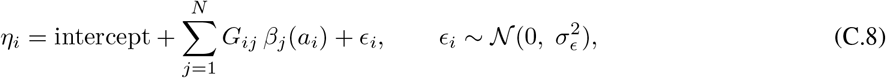

and the binary label is *y*_*i*_ *~* Bernoulli(*σ*(*η*_*i*_)).

#### C.5 Control Knobs and Stress-Test Axes

The two experimental axes in the main paper correspond directly to HGP parameters. *Effect-size non-stationarity* is controlled by the lengthscale multiplier *µ*_*s*_: tasks with *µ*_*s*_ = 0.008 exhibit strong local drift, while tasks with *µ*_*s*_ = 100.0 are nearly stationary. *Training-set dispersion* is controlled by the cluster width multipliers and the majority-cluster dominance ratio: increasing the width of minority clusters or reducing the majority-to-minority ratio expands the ancestry spread of the sampled population.

For instruction tuning, the overall population drawn from the HGP generator remains multi-ancestry, but the labeled support set presented to the model is sampled with a majority-cluster skew. This forces the model to learn to extrapolate from an imbalanced context to a more diverse query set, directly mimicking the failure mode identified in the main paper.

### D PRS-CSx-Inspired Task Family

The second task family adapts the continuous-shrinkage logic of PRS-CSx (Ruan et al., 2022) to a continuous ancestry manifold. The goal is to preserve biologically meaningful ingredients that are absent from a pure GP generator: sparse heavy-tailed effects, ancestry-specific allele frequencies, cluster-specific heritability, and imperfect cross-population effect correlation.

#### D.1 Global-local effect prior

Let *z*_*j*_ *∈* {0, 1} denote whether SNP *j* is causal. For each SNP we sample a global-local shrinkage hierarchy

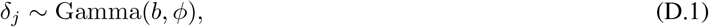

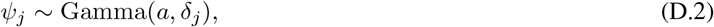

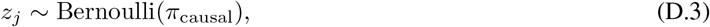

where *φ* controls overall sparsity and *ψ*_*j*_ determines the local scale of SNP *j*. This produces a heavy-tailed prior with strong shrinkage around zero and a small number of large effects.

For each causal SNP, population-level effects are sampled jointly across ancestry clusters:

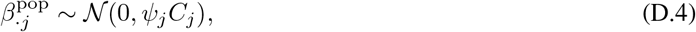

where *C*_*j*_ is a cross-population covariance matrix. Its diagonal entries encode per-cluster effect scale and its off-diagonal entries are governed by a per-SNP genetic-correlation parameter *r*_*g,j*_. This lets some loci be nearly shared across ancestry clusters while others diverge appreciably.

#### D.2 Allele frequencies and MAF-dependent architecture

The PRS-CSx family uses a Balding–Nichols construction for ancestry-specific allele frequencies. First an ancestral minor-allele frequency is sampled as

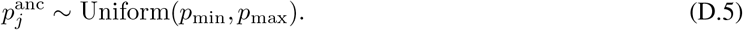

Then each cluster-specific frequency is sampled from a Beta distribution whose variance is controlled by cluster-specific *F*_*ST*:_

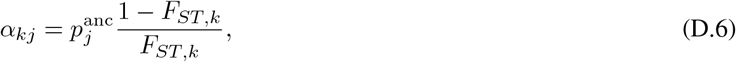

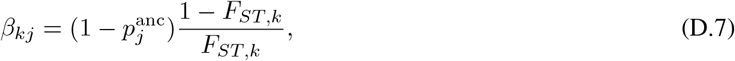

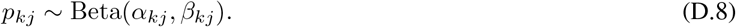

Genotype dosages are then sampled as *G*_*ij*_ *~* Binomial 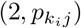 for subjects assigned to cluster *k*_*i*_.

To model the empirical tendency of rarer variants to carry larger per-allele effects, we optionally rescale cluster-level effect sizes by allele frequency,

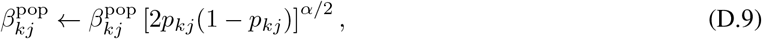

with *α* = 0 recovering no rescaling and *α <* 0 amplifying low-frequency variants. Because *p*_*kj*_ differs across ancestry clusters, this mechanism itself contributes to non-stationarity.

#### D.3 Continuous ancestry interpolation and phenotype model

PRS-CSx is population-based, so we extend it to a continuous ancestry coordinate by interpolating cluster-level effects. Each SNP receives its own interpolation bandwidth *h*_*j*_ sampled from a task-specific range. For subject *i*, cluster *k*, and SNP *j*, unnormalized weights are

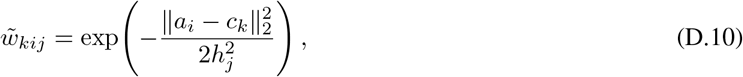

and normalized weights satisfy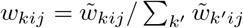. The subject-specific effect for SNP *j* is then

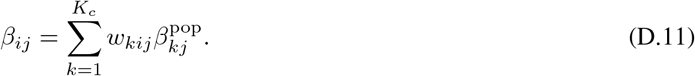

Given genotype matrix *G*, the genetic value is *g*_*i*_ = ∑ ^*j*^ *G*_*ij*_*β*_*ij*_. Residual noise is calibrated by a target cluster-specific heritability 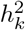 through

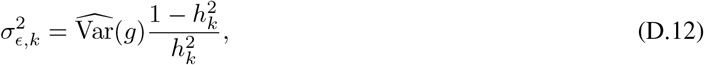

where 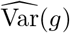 is the empirical population-wide variance of the genetic values. The latent liability is *η*_*i*_ = *g*_*i*_ + _*i*_ with 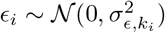, and the final label is obtained after logistic transformation and per-cluster thresholding. Per-cluster thresholding keeps prevalence approximately balanced within each ancestry cluster and prevents a model from exploiting simple ancestry-prevalence shortcuts.

#### D.4 Why PRS-CSx is more biologically grounded than HGP

The HGP family is useful precisely because it is analytically transparent, but the PRS-CSx family is closer to the structure of real genomic prediction tasks. The differences are substantive. HGP treats effect functions as smooth Gaussian draws over ancestry, whereas PRS-CSx imposes sparse heavy-tailed effects with explicit global-local shrinkage. HGP typically treats genotype generation and effect variation as loosely coupled, whereas PRS-CSx connects effect magnitudes to cluster-specific allele frequencies, heritability, and cross-population genetic correlation. Finally, PRS-CSx lets different loci vary across ancestry at different spatial scales through SNP-specific interpolation bandwidths. These ingredients make PRS-CSx a better proxy for biologically structured non-stationarity, even though it remains synthetic.

### E Mixed Instruction-Tuning Protocol

#### E.1 Data and Episode Construction

The mixed generator creates 400 training settings and 90 held-out evaluation settings, drawn approximately uniformly from the HGP and PRS-CSx families. Each accepted setting produces a train pickle (at least 5,000 individuals from the training ancestry distribution), a test pickle (at least 100 individuals from the shifted test ancestry distribution), and a latents file recording ancestry coordinates, genotypes, effect sizes, cluster identities, and setting metadata. The two families share the same tabular schema, so instruction tuning can sample episodes from both families without changing the model input format. A manifest CSV indexes the settings; one episode (one in-context learning task) is built from one manifest row by sampling *N*_tst_ = 0.9 × |test test rows as the masked query and *N*_ctx_ = 10 × |*N*_tst_ rows from the train pickle as labeled context. Features are zero-padded to a common feature dimension and labels are remapped to contiguous integers. The model receives the concatenated sequence [context | query] in a single forward pass.

#### E.2 Stage-3 Fine-Tuning Only

TabICL is composed of three modules: a column embedder, a row interactor, and an in-context learning head (Qu et al., 2025). We perform Stage-3 fine-tuning only: the column embedder and row interactor are frozen at their public-checkpoint weights, and gradient updates are applied exclusively to the in-context learning head. This preserves the general-purpose feature- and row-level representations learned during the upstream pretraining stages while specializing the in-context reasoning head to ancestry-shifted genotype-to-phenotype tasks.

#### E.3 Loss and Optimizer

The training loss is per-token cross-entropy on the masked query positions only,

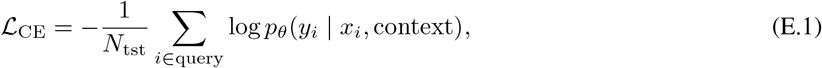

augmented with a proximal anchor regularizer toward the initial off-the-shelf checkpoint *θ*_0_,

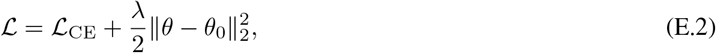

where *θ* collects the trainable parameters of the in-context learning head and *λ* is the swept proximal coefficient (interpreted as a weight-decay value in the HPO grid). This proximal term keeps the tuned head close to the pretrained initialization, mitigating catastrophic forgetting of the pretraining distribution while the model adapts to ancestry-shifted tasks. Optimization uses AdamW with cosine learning-rate decay and a 10% linear warmup, gradient clipping at norm 1.0, and float32 precision.

#### E.4 Per-Step Update Mechanics

A single gradient update consumes a batch of *B* = 10 episodes. To accommodate variable per-task sequence lengths and reduce peak GPU memory, each batch is split into *B/B*_*µ*_ = 10 micro-batches of size *B*_*µ*_ = 1. For each micro-batch the forward pass produces query-token logits, the cross-entropy loss is scaled by 1*/B* and back-propagated, and gradients are accumulated. After all 10 micro-batches the AdamW optimizer takes a single step and gradients are zeroed:

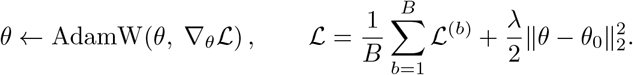

Each run trains for 500 such gradient updates.

#### E.5 Task Switching

Tasks are switched at the episode level: each of the *B* = 10 episodes inside a gradient update is an independent random draw (with replacement) from the 400-setting training manifest, so a single weight update sees gradient signal from up to ten distinct synthetic task settings sampled from both the HGP and PRS-CSx families. Because successive gradient steps redraw fresh manifest rows, the sequence of training tasks is i.i.d. across the manifest and not a fixed curriculum. Within an episode the support and query rows are themselves resampled (without replacement on the test side, with replacement when more than |train| */*10 context rows are needed), so the same setting yields a new episode each time it is drawn.

#### E.6 Per-Step Evaluation and Best-Checkpoint Selection

Before the first weight update, a fixed Stage-1 evaluation batch is constructed once from the first 30 rows of the held-out evaluation manifest using a fixed seed, so that the same 30 episodes are reused at every step and metrics are directly comparable across steps. After each gradient update we compute, on this fixed batch, the mean cross-entropy and the mean hard-label accuracy (the fraction of query tokens whose argmax prediction matches the true label). Whenever the running eval accuracy exceeds the previous best, the current parameters are written to disk as the run’s best checkpoint and the previous best is deleted; periodic checkpoints are also saved every 5 steps for diagnostic purposes.

#### E.7 HPO Search and Best-Setting Selection

The instruction-tuning hyperparameters are selected by a grid search over learning rate and proximal weight-decay coefficient,

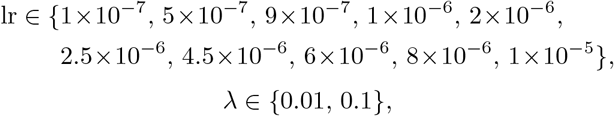

yielding 20 runs that share all other configuration. After every run completes, an unbiased Stage-2 evaluation is performed on a separate slice of the evaluation manifest (rows 30–64) that the training loop never observed. Five checkpoints are scored per run—the best Stage-1 checkpoint plus the four periodic checkpoints whose step indices are closest to it—and the per-run Stage-2 score is the median accuracy and median cross-entropy across these five checkpoints, which damps step-to-step noise in the HPO ranking. The HPO winner is the run with the highest median Stage-2 accuracy; for the sweep used in the main paper this corresponds to lr = 4.5×10^−6^ and *λ* = 0.1. Figures E.1 and E.2 show the full sweep on a lr × *λ* grid.

#### E.8 Compute Resources

Instruction tuning and hyperparameter search were run on a university-managed SLURM cluster. The instruction-tuning jobs used one or two NVIDIA V100 GPUs with 32 GB memory per GPU, depending on scheduling availability and the specific job stage. For the HPO sweep reported here, the full university-cluster run completed in less than 24 hours wall-clock time. The downstream AoU evaluations were run in the Google Cloud environment through the AoU Researcher Workbench, using an NVIDIA V100 GPU with 16 GB memory. This resource is provided directly by AoU on a pay-per-use basis at approximately 3 USD per hour, and the full set of evaluations reported in the paper required less than 3 hours of runtime.

#### E.9 Downstream Use

The downstream real-data evaluation in the main paper uses no additional fine-tuning on AoU. Both the off-the-shelf and tuned models operate in the same in-context regime, so the comparison isolates the effect of synthetic-task alignment rather than real-data parameter updates.

### F Asset Release and Responsible Use

The new assets contributed by this project are the synthetic task generators, the mixed task manifests, and the instruction-tuned TabICL checkpoint. These assets can be released without exposing individual-level AoU data because all downstream controlled-tier information remains outside the released artifacts. Since the work concerns genomic prediction, release materials should explicitly state that ancestry coordinates are analytic summaries of genetic variation, not social identity labels, and that the tuned model is intended for controlled research use rather than clinical deployment. The main paper discusses the fairness motivation for the work; this appendix clarifies that release is limited to synthetic artifacts and model weights.

## Acknowledgements

This work builds on off-the-shelf models and software released by the teams behind *TabICL: A Tabular Foundation Model for In-Context Learning on Large Data* (Qu et al., 2025) *and TabPFN. We gratefully acknowledge the researchers and developers who created and released these resources. In particular, our experiments use the public off-the-shelf TabICL checkpoint (version 0*.*1*.*4) and the public off-the-shelf TabPFN release (version 6*.*3*.*2), with no task-specific retraining for the off-the-shelf evaluations. We also gratefully acknowledge All of Us participants for their contributions, without whom this research would not have been possible, and we thank the National Institutes of Health’s All of Us Research Program for making available the participant and sample data examined in this study. All analyses were conducted on controlled-access data; no participant-level genetic or phenotype data are reported, and all results shown in the paper are aggregated over cohorts larger than 50 participants to reduce disclosure risk. In preparing the manuscript, we used agentic language models, including Claude and GPT, to help correct grammar and spelling, to improve the presentation of the writing, and to write code; all technical content, AI-generated code, and final wording decisions were reviewed by the authors*.

---

**Figure B.1.**
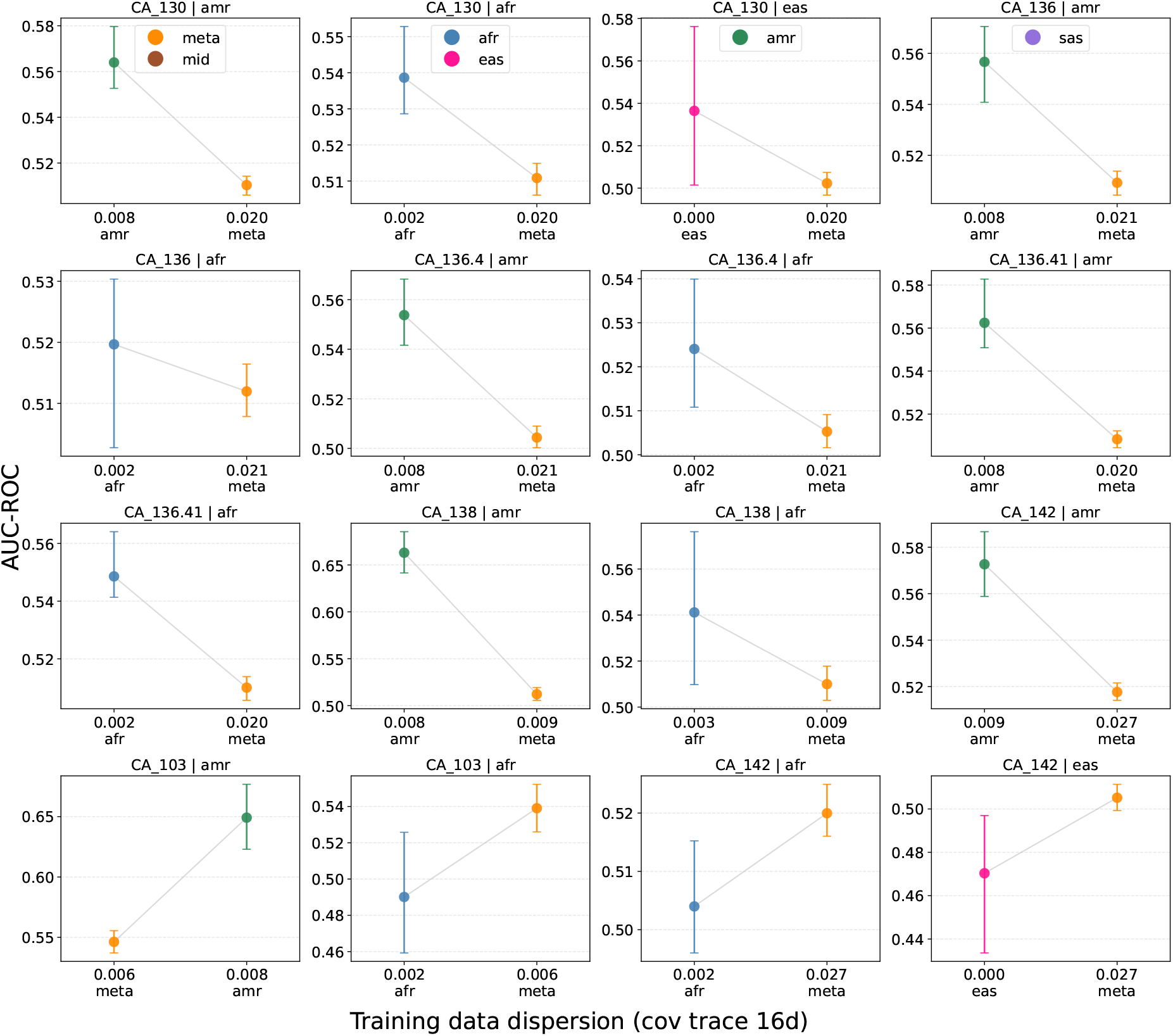
Matched-dispersion real-data analysis for off-the-shelf TabPFN version 6.3.2 on AoU cancer phenotypes. Each panel fixes a phenotype and target ancestry, then compares an ancestry-specific version 7 context table against a size-matched meta-ancestry version 7 context table. Evaluation uses held-out version 8 samples. In 12 of 16 matched comparisons, TabPFN performs better with the less dispersed ancestry-specific context table; intervals show 60% bootstrap confidence intervals.

**Table B.1:** Phenotype-code mapping for the cancer and respiratory AoU evaluations. The figures in the main paper use phenotype codes for compactness; this table gives the corresponding phenotype names and the number of EUR version 7 training cases available for each phenotype.

| Family | Code | Full phenotype name | EUR v7 cases |
| --- | --- | --- | --- |
| Cancer | CA_103 | Malignant neoplasm of the skin | 10,802 |
| Cancer | CA_130 | Cancer (solid tumor, excluding BCC) | 14,392 |
| Cancer | CA_136 | Benign neoplasm of the digestive organs | 12,174 |
| Cancer | CA_136.4 | Benign neoplasm of colon, rectum, anus and anal canal | 11,521 |
| Cancer | CA_136.41 | Benign neoplasm of the colon | 11,260 |
| Cancer | CA_138 | Benign neoplasm of the skin | 13,328 |
| Cancer | CA_142 | Lump or mass in breast or nonspecific abnormal breast exam | 10,124 |
| Respiratory | RE_460 | Respiratory infection | 19,385 |
| Respiratory | RE_460.1 | Acute upper respiratory infection | 16,645 |
| Respiratory | RE_462 | Sinusitis | 11,675 |
| Respiratory | RE_463 | Rhinitis and nasal congestion | 14,683 |
| Respiratory | RE_463.2 | Allergic rhinitis | 11,603 |
| Respiratory | RE_475 | Asthma | 10,566 |
| Respiratory | RE_488 | Abnormalities of breathing | 20,690 |
| Respiratory | RE_488.1 | Dyspnea [Shortness of breath] | 15,486 |
| Respiratory | RE_499 | Other symptoms and disorders of the respiratory system | 17,137 |
| Respiratory | RE_499.1 | Cough | 15,966 |

**Figure E.1.**
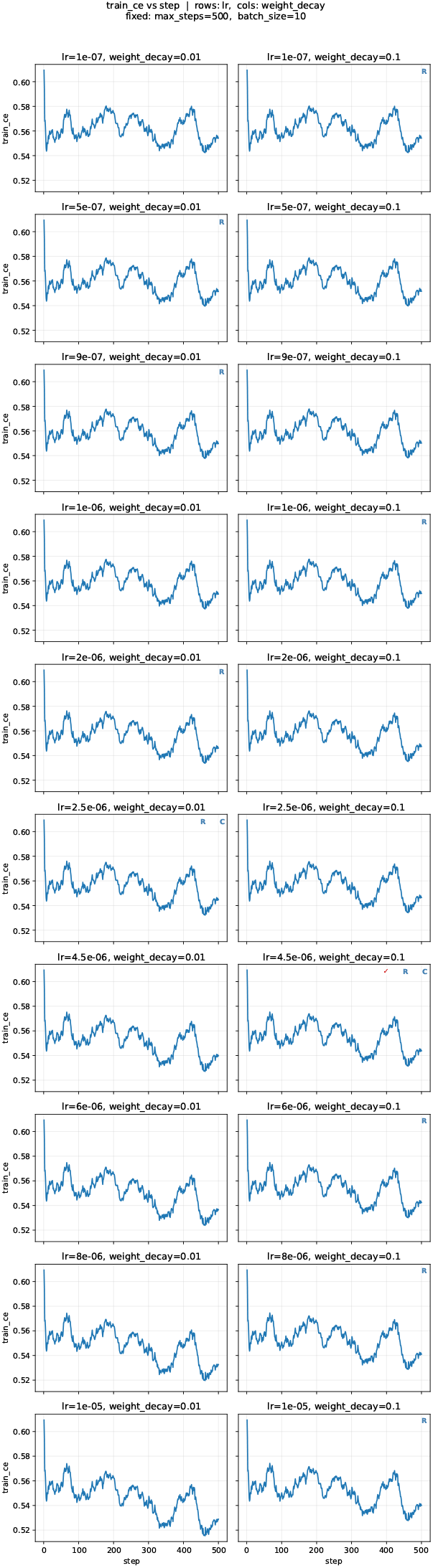
Per-step training cross-entropy across the instruction-tuning HPO grid. Rows index learning rate (10 values from 10^−7^ to 10^−5^), columns index the proximal weight-decay coefficient *λ ∈* {0.01, 0.1}. Each subplot shows training cross-entropy as a function of gradient-update step for that single (lr, *λ*) cell.

**Figure E.2.**
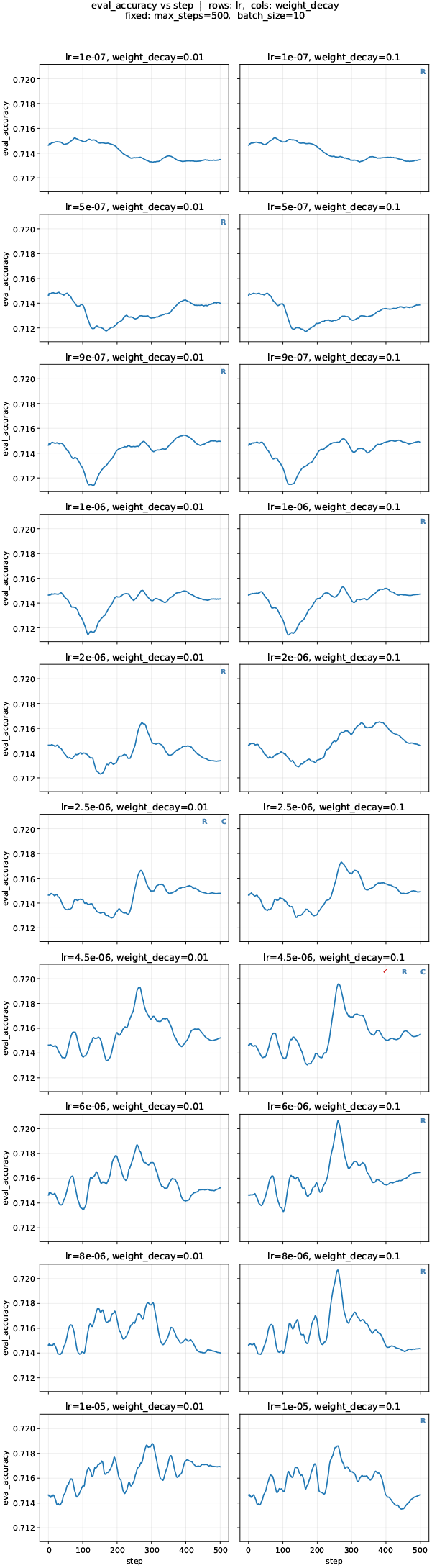
Per-step held-out Stage-1 evaluation accuracy across the same HPO grid. Layout matches Figure E.1: rows are learning rates and columns are proximal weight-decay coefficients. Each subplot shows mean hard-label accuracy on the fixed 30-task Stage-1 evaluation batch, reused at every step. The best Stage-2 cells are highlighted; the global winner is lr = 4.5×10^−6^, *λ* = 0.1.

## References

All of Us Research Program. How the all of us genomic data are organized. All of Us Research Program support page, 2024.

All of Us Research Program Genomics Investigators. Genomic data in the all of us research program. Nature, 627 (8003):340–346, 2024.

S. A. Bien, G. L. Wojcik, C. J. Hodonsky, et al. The future of genomic studies must be globally representative: Perspectives from page. Annual Review of Genomics and Human Genetics, 20:181–200, 2019.

S. W. Choi, T. S.-H. Mak, and P. F. O’Reilly. Tutorial: a guide to performing polygenic risk score analyses. Nature Protocols, 15(9):2759–2772, 2020.

A. Das and Y. Cui. Bridging ancestry gaps in genomic risk prediction with tabular foundation models. Bioinformatics, 2026. Vol. 42, Supplement_1, article btag217. DOI: 10.1093/bioinformatics/btag217.

A. Das, M. I. Khalid, R. Peñaloza, et al. When no paths lead to Rome: Benchmarking systematic neural relational reasoning. In Advances in Neural Information Processing Systems (Datasets and Benchmarks Track), volume 38, 2025.

A. Das, J. Boisson, I. Khalid, et al. Project Auto-World: Towards automated benchmarking of neural relational reasoners. arXiv preprint arXiv:2606.24965, 2026.

Y. Ding, K. Hou, Z. Xu, et al. Polygenic scoring accuracy varies across the genetic ancestry continuum. Nature, 618: 774–781, 2023.

Y. Gao and Y. Cui. Deep transfer learning for reducing health care disparities arising from biomedical data inequality. Nature Communications, 11(1):5131, 2020.

D. Gurdasani, I. Barroso, E. Zeggini, et al. Genomics of disease risk in globally diverse populations. Nature Reviews Genetics, 20:520–535, 2019.

K. Helli, D. Schnurr, N. Hollmann, et al. Drift-resilient TabPFN: In-context learning temporal distribution shifts on tabular data. In Advances in Neural Information Processing Systems, 2024.

N. Hollmann, S. Muller, L. Purucker, et al. Accurate predictions on small data with a tabular foundation model. Nature, 637:319–326, 2025.

L. Kachuri, N. Chatterjee, J. Hirbo, et al. Principles and methods for transferring polygenic risk scores across global populations. Nature Reviews Genetics, 25(1):8–25, 2024. doi: 10.1038/s41576-023-00637-2.

P. C. Mahalanobis. On the generalized distance in statistics. Sankhyā: The Indian Journal of Statistics, Series A (2008-), 80:S1–S7, 2018.

M. C. Mills and C. Rahal. The gwas diversity monitor tracks diversity by disease in real time. Nature Genetics, 52: 242–243, 2020.

R. E. Peterson, K. Kuchenbaecker, R. K. Walters, et al. Genome-wide association studies in ancestrally diverse populations. Cell, 179(3):589–603, 2019.

J. Qu, D. Holzmüller, G. Varoquaux, et al. TabICL: A tabular foundation model for in-context learning on large data. In International Conference on Machine Learning, pages 50817–50847, 2025.

Y. Ruan, Y.-F. Lin, Y.-C. A. Feng, et al. Improving polygenic prediction in ancestrally diverse populations. Nature Genetics, 54(5):573–580, 2022.

M. Seeger. Gaussian processes for machine learning. International Journal of Neural Systems, 14(2):69–106, 2004.

G. Sirugo, S. M. Williams, and S. A. Tishkoff. The missing diversity in human genetic studies. Cell, 177(1):26–31, 2019.

C. Tcheandjieu, X. Zhu, A. T. Hilliard, et al. Large-scale genome-wide association study of coronary artery disease in genetically diverse populations. Nature Medicine, 28(8):1679–1692, 2022.

V. Thomas, J. Ma, R. Hosseinzadeh, et al. Retrieval & fine-tuning for in-context tabular models. In Advances in Neural Information Processing Systems, volume 37, pages 108439–108467, 2024.

Y. Wang, M. Kanai, T. Tan, et al. Polygenic prediction across populations is influenced by ancestry, genetic architecture, and methodology. Cell Genomics, 3:100408, 2023.

J. Wei, M. Bosma, V. Y. Zhao, et al. Finetuned language models are zero-shot learners. arXiv preprint arXiv:2109.01652, 2021.

